# Synchronized oscillations, metachronal waves, and jammed clusters in sterically interacting active filament arrays

**DOI:** 10.1101/2020.06.08.140731

**Authors:** Raghunath Chelakkot, Michael F. Hagan, L. Mahadevan, Arvind Gopinath

## Abstract

Autonomous active, elastic filaments that interact with each other to achieve cooperation and synchrony underlie many critical functions in biology. A striking example is ciliary arrays in the mammalian respiratory tract; here individual filaments communicate through direct interactions and through the surrounding fluid to generate metachronal traveling waves crucial for mucociliary clearance. The mechanisms underlying this collective response and the essential ingredients for stable synchronization remain a mystery. In this article, we describe Brownian dynamics simulations of multi-filament arrays, demonstrating that short-range steric inter-filament interactions and surface-roughness are sufficient to generate a rich variety of collective spatiotemporal oscillatory, traveling and static patterns. Starting from results for the collective dynamics of two- and three-filament systems, we identify parameter ranges in which inter-filament interactions lead to synchronized oscillations. We then study how these results generalize to large one-dimensional arrays of many interacting filaments. The phase space characterizing the multi-filament array dynamics and deformations exhibits rich behaviors, including oscillations and traveling metachronal waves, depending on the interplay between geometric spacing between filaments, activity, and elasticity of the filaments. Interestingly, the existence of metachronal waves is nonmonotonic with respect to the inter-filament spacing. We also find that the degree of filament surface roughness significantly affects the dynamics — roughness on scales comparable to the filament thickness generates a locking-mechanism that transforms traveling wave patterns into statically stuck and jammed configurations. Our simulations suggest that short-ranged steric inter-filament interactions are sufficient and perhaps even critical for the development, stability and regulation of collective patterns.

## 1 Introduction

The emergence of oscillations in single or arrayed elastic filamentous structures, such as the graceful rhythmic movements of ciliary beds, is a common motif in biology^1–5^. In these systems, emergent collective oscillations and waves are strongly affected by multiple effects, including the elasticity of the underlying filamentous structures, modes of activation due to molecular motors, coupling between neighboring filaments, and boundaries^6–16^. Due to the complexity and many-body nature of these systems, disentangling the contributions of each of these effects to the system dynamics is highly challenging.

Inspired by the manner in which these biological active filamentous carpets integrate elasticity with biological motor activity to generate sustained and stable oscillations, a number of reconstituted or synthetic *active filament* systems have been developed^17–28^. Here activity is imbued either by motors acting externally on filaments to generate elastic forces^17–19^, or by using internally propelled filaments constructed of beads that are powered by surface chemical reactions or responses to external fields. Within this class of systems, the interplay between geometry, filament elasticity, and activity can result in non-linear buckling instabilities, where initially stationary filaments yield to stable oscillations and periodic motions. These instabilities have been the subject of recent theoretical and computational inquiries. Continuum as well as discrete agent-based models have been used to investigate the emergence of oscillations in single filaments, and coupling-induced synchrony in systems of two rotating filaments^29–40^.

However, an equally important set of problems — the collective behaviors of many elastic active filaments — have yet to be investigated in detail. In this case, key questions are as follows: first, how do individual filaments - each capable of beating autonomously - respond to the force fields generated by their neighbors and synchronize their spatiotemporal patterns? In particular, under what conditions do such systems exhibit stationary states characterized by propagating spatiotemporal patterns and waves? Second, how are the properties of filament waveforms regulated by the geometric, elastic and active aspects of the collective system?

In these low Reynolds number, actively driven, collective systems, two types of inter-filament interactions are expected to play a crucial role (in addition to single filament properties such as elasticity and activity). The first type of inter-filament interaction comprises fluid-mediated medium and long-range elasto-hydrodynamic interactions41 that alter the viscous drag on filaments and couple to their spatiotemporal response. A recent analytical study^33^ by one of us analyzed the onset of synchronization of clusters, arrays and carpets of active filaments grafted to a rigid impenetrable planar wall. Full multi-filament and filament-wall hydrodynamic interactions were considered using singularity methods built on slender body theory. Stable, oscillatory states where filaments oscillated with the same frequency with a varying phase angle were determined to bifurcate from a stationary state. Similar computational studies and phenomenological models that used the simpler resistivity approximation to treat fluid mediated interactions^42–46^ have also demonstrated that hydrodynamic interactions can lead to stable collective and synchronized responses.

The second type of interaction includes short-range effects, such as steric interactions, electrostatic interactions and surfaceroughness-based frictional effects. Recent studies suggest that steric interactions and collective fluid mechanical effects both play important roles in biologically relevant multi-filament arrays, such as in the passive arrayed brush-like structures in the glycocalyx^41^ and active ciliary carpets in the mucociliary tract^47^. Also clear from these studies is the important role of surface-attached features and networked structures. For instance, Button *et. al*.^47^ proposed a Gel-on-Brush model of the mucus clearance system, in which the periciliary layer is occupied by membrane-spanning mucins and large mucopolysaccharides that are tethered to cilia and microvilli. They hypothesize that the tethered macromolecules produce inter-molecular repulsions, which stabilize the layer against compression by an osmotically active mucus layer.

Here, we complement previous studies that neglect steric, contact and roughness interactions, by performing Brownian Dynamics (BD) simulations with these effects included. Motivated by observations on the brush-like structure of cilia^47^, we consider each filament to interact with its neighbors via a steric potential with an interaction length-scale ***σ*** that is comparable to, but may differ from, the intrinsic geometric filament thickness - that is the segment length *ℓ*_0_. Adjusting the ratio of these two scales allows us to vary the inter-filament interactions between the regimes of *smooth* (***σ*** > *ℓ*_0_) or *rough* filaments (***σ*** = *ℓ*_0_). Thus, we are able to study both the effect of excluded volume and surface roughness on collective behaviors of active filament arrays.

The layout of the article is as follows. We introduce our computational model for an active filament system in §2; the system consists of a small cluster (2-3 filaments) or a large periodic array (300 filaments) immersed in a viscous constant temperature fluid (Figure 1(a)). Each filament comprises connected active beads and is geometrically fixed at one end. The other end is free, and this degree of freedom allows each filament to independently and autonomously oscillate in a planar manner via active buckling instabilities. The intrinsic frequency and amplitude of beating by individual filaments is controlled by the interplay between the filament geometry, elasticity, fluid dissipation, and activity. In the absence of inter-filament interactions, adjacent filaments beat with the same frequency but are generally out of phase. In §3, we analyze the collective dynamics and emergent steric-driven coupling in small clusters comprising 2 or 3 smooth filaments. Building on this, we analyze the dynamics of large arrays comprised of smooth filaments in §4. We next probe the effect of surface corrugation within the framework of the steric model introduced in §2, by setting ***σ*** ≈ *ℓ*_0_, resulting in large gradients of the excluded-volume potential between adjacent filaments. Effectively, beads in neighboring filaments interlock as they move, resulting in higher effective friction coefficients and significantly reducing their tangential velocities. This extra friction results in qualitatively different collective dynamics in comparison to the smooth filaments. The final set of results explores relaxing the hard constraint (clamped base) by implementing a softer constraint (pivoting base). We conclude in §6 and highlight features that are relevant to previous studies and serve as motivation for future experimental and computational work.

**Fig. 1.**
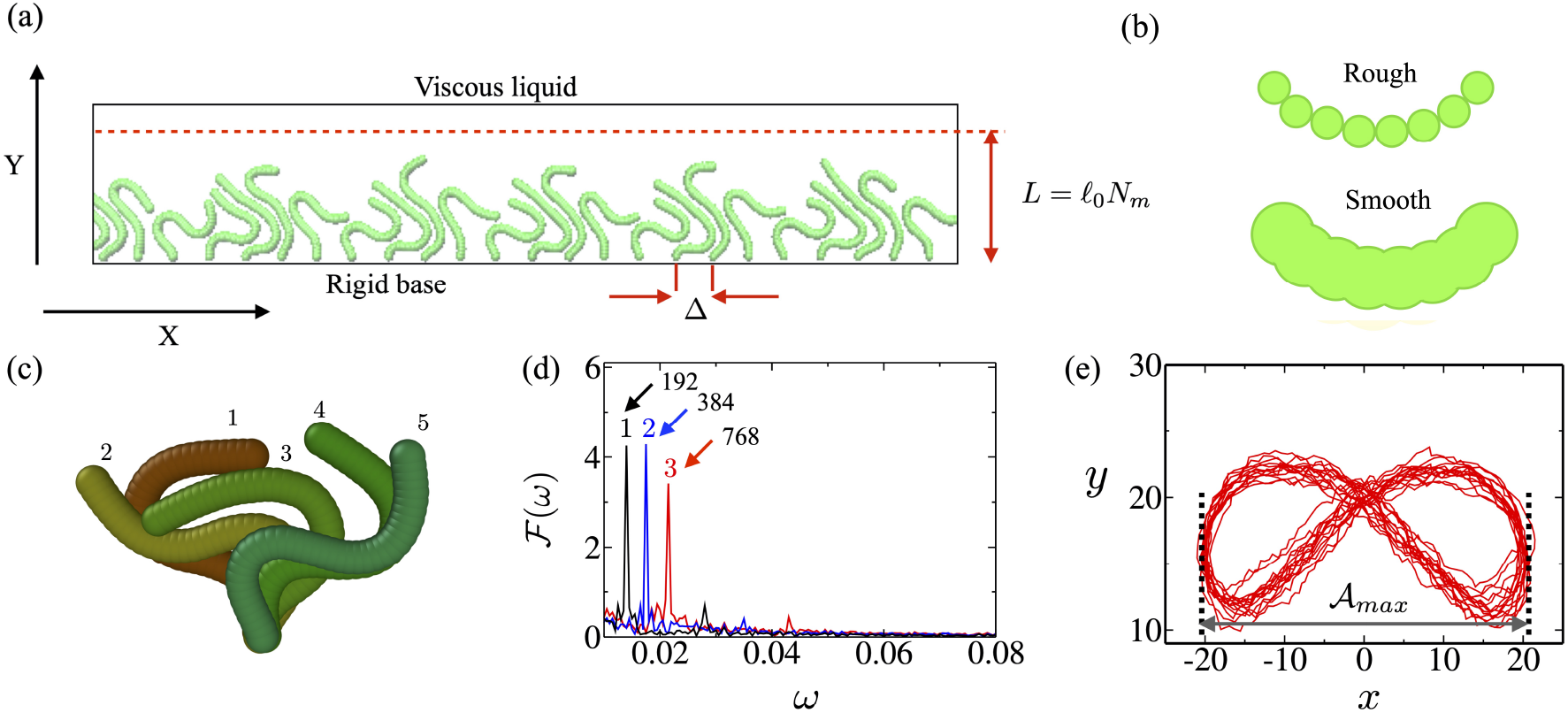
(a) Typical arrangement of clamped, active filaments in the 1D array used in the simulations. Each active filament is comprised of *N*_m_ connected spherical self-propelling beads (discs), with each pair separated by distance *ℓ*_0_ in the undeformed state. (b) Schematic of rough (upper image) and smooth (lower image) filament structures. Both have the same bead size *ℓ*_0_; however the smooth filament has a larger effective steric interaction lengthscale *σ* > *ℓ*_0_. This mimics a brush filament as in the mucociliary tract^47^. The intrinsic elasticity of the filaments (set by the parameters *B* and *K*_E_) are the same in both cases; however, the overlapping spheres make the effective surface of the smooth filament less corrugated than that of its rough counterpart. (c) Typical configurations of an isolated filament during an oscillatory cycle (indices 1-5 represent times) for dimensionless activity strength 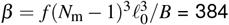. Note that *ℓ*_0_ = 1 in reduced (simulation) units. (d) Fourier transform 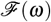 of the end-end distance *L*_ee_ indicating distinct frequency peaks at (1) *β* = 192, (2) *β* = 384, and (3) *β* = 768. (e) The trajectory of the end-segment of a filament with *β* = 192. 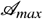 denotes the maximum displacement of the end-segment along the *x* direction, averaged over many oscillatory cycles.

Our studies reveal that short-range repulsive interactions can alter the oscillation phase of filaments to generate global synchronization. The phase-space characterizing the dynamical response of the array exhibits rich behaviors, with oscillations and traveling waves appearing, vanishing, and eventually re-appearing depending on the parameter values. We also show that roughness at the filament scale yields an additional locking-mechanism that dramatically changes the form and wave-speed of metachronal waves. Finally, relaxing the strength of the geometric constraint at the base and allowing for flexible pivoting results in jammed static shapes, even though the system itself remains active and dynamic.

## 2 Computational Model

The active filament carpet/array comprises *N* two-dimensional active filaments (chains) arrayed uniformly in one dimension along the *e_x_* direction and initially aligned along the *e_y_* direction as illustrated in Figure 1(a). The spacing between the filaments, Δ, is treated as an adjustable parameter in the simulations. We consider sparse carpets comprised of only a few filaments (*N* = 2 and *N* = 3) and then a larger carpet with (*N* = 300) more filaments.

To focus on the role of steric interactions, we do not consider hydrodynamic interactions and the wall only serves to keep the base of the filament fixed. Note that in a system where full hydrodynamic effects are included, fluid flow generated by beating filaments will alter the motion of the filament^33–35^. In our case, we neglect these induced fluid flows and consider, to leading order, just the viscous Stokes drag on the beads comprising the filament as they move.

In the following we introduce potentials that are used to calculate extensional, bending, and steric forces. Dimensional potentials are starred; all potentials are scaled with *k*_B_*T* with *T* the thermodynamic temperature of the ambient fluid.

### 2.1 Interaction potentials

Each active filament is a collection of *N*_m_ polar, active spheres (disks) of effective diameter ***σ*** in 2D as shown in Figure 1(b). The coordinate of the **α**^th^ sphere is denoted by **r**_α_ and it is connected to the neighbouring spheres of the same filament via extensional and bending potentials as illustrated in Figure 1(c)).

The extensional force between adjacent beads is derived from the total potential 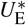 given by

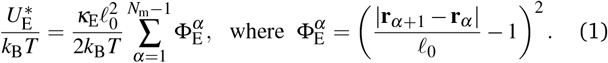

The value of κ_E_ is maintained at a value large enough that the actual distance between each polar particle is nearly *ℓ*_0_, making the chain nearly inextensible. The overall length of the undeformed filament is thus *ℓ* = (*N*_m_ – 1)*ℓ*_0_.

The overall resistance to bending is implemented via a three-body bending potential motivated by the energy for a thin elastic continuous curve in the noise-less limit,

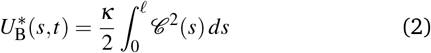

where 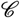 is the curvature measured along the centerline of the curve. We discretize (2) for our model filaments by approximating the curvature at bead *α* using 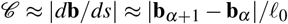, where **b**_*α*_ = (**r**_*α*−1_ − **r**_*α*_)/|**r**_*α*_ − **r**_*α*−1_| is the unit bond vector that is anti-parallel to the local tangent.

In the continuous limit (*ℓ*_0_ → 0, *N*_m_ → ∞, *N*_m_*ℓ*_0_ → constant), **b**_*α*_ identifies with the tangent vector **t** of the continuous model at arclength *s* = α*ℓ*_0_; thus (**b**_*α*+1_ − **b**_*α*_)/***σ*** ≈ *d***t**/*ds*. Discretizing (2) using *B* ≡ *κ*/*ℓ*_0_, we write

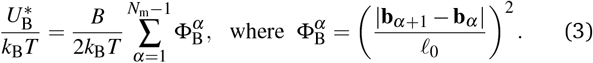

We account for excluded-volume (steric) interactions between beads in neighboring filaments via a short-range repulsive WCA (Weeks-Chandler-Anderson) interaction potential. Here, we have chosen filament lengths and rigidity values such that overlap between beads in the same element does not occur. With *r_αβ_* ≡ |**r**_*α*_ − **r**_*β*_| as the distance between a pair of spheres (*α, β*) belonging to different filaments, the net overall steric potential summed over all segments (beads) is

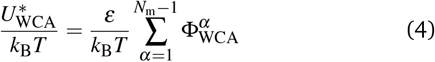

where

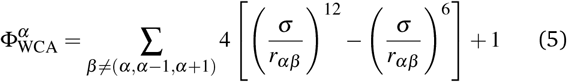

if 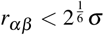 and *u*(*r*) = 0 otherwise. The index *β* refers to pairs of beads in the same filament as well as in neighboring filaments, thus incorporating all possible steric interactions. In (4), *ε* = *k_B_T*.

The effect of the steric interactions encoded in the interaction potentials (4) and (5) depends on the softness of the interaction potential and also on the fine structure and roughness of the interacting filament. The former effect is controlled by the power-law exponents in the WCA, while the latter can be varied by changing the ratio *ℓ*_0_/***σ***. Thus the length-scale ***σ*** effectively sets the nature and the scale of the steric excluded volume interactions.

Each disc comprising the filament is self-propelling with a velocity *v*_0_**b**_*α*_, in the direction of the local tangent **b**_*α*_ of the filament. This causes local compression, generating *follower* forces of magnitude *F* that follow the local target of the filament. In the continuous and over-damped limit, this yields a uniform active force per unit length. Since *v*_0_ is a constant for each bead on the filament, the quantity *v*_0_ = *μF* is also constant for each realization and can be interpreted as the magnitude of the active force exerted by each bead. We note that the magnitude of the total force for a straight unbent filament ~ *v*_0_*N*_m_/*μ*, so the effective force density *f* = *F*/*ℓ*_0_ ~ (*v*_0_*N*_m_/*μ*)/(*N*_m_*ℓ*_0_).

### 2.2 Equations of motion

We evolve the position **r**_*α*_ of each bead *α* using Brownian dynamics, with the forces accounting for extensional, bending, steric, and thermal effects described above. We render equations dimensionless by scaling quantities as follows. We use *ℓ*_0_ as the unit of length, the diffusive relaxation time 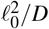 as unit of time, and *k_B_T* as the unit of energy. In the over-damped limit, the equations of motion can be written as

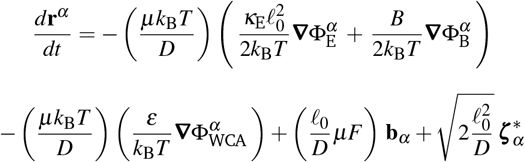

Here 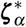 is a delta-correlated noise with zero mean acting on the disc. With the units of length, time, and energy defined above, the mobility *μ* = *D*/*k_B_T* = 1 in dimensionless form. Other parameters in the dimensionless (reduced) units are listed in Table 1. The equations of motion in dimensionless form then reduce to

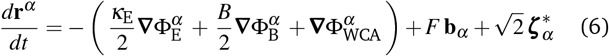

**Table 1.**
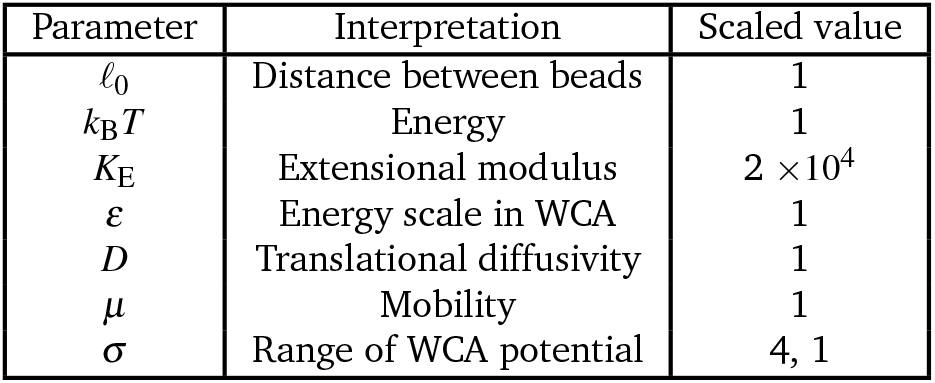
List of parameters held constant in simulation and their value in scaled units

| Parameter | Interpretation | Scaled value |
| --- | --- | --- |
| $\ell_0$ | Distance between beads | 1 |
| $k_B T$ | Energy | 1 |
| $K_E$ | Extensional modulus | $2 \times 10^4$ |
| $\varepsilon$ | Energy scale in WCA | 1 |
| $D$ | Translational diffusivity | 1 |
| $\mu$ | Mobility | 1 |
| $\sigma$ | Range of WCA potential | 4, 1 |

Interpreting the time derivative in the Ito-Stratanovich sense, we solve Eq. (6) using a time-stepper based on the Euler-Maruyama scheme. Theory29 shows that the behavior of an isolated active filament depends on a single effective *activity parameter β* ≡ *fℓ*/*κ*. In our case the *force density f* is related to the force on a bead *F* by *f* = *F*/*ℓ*_0_, so that

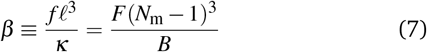

in simulation units.

### 2.3 Simulation conditions and parameters

We present simulations for two limiting cases in §3. The first set considers *smooth filaments*, with scaled value ***σ*** = 4; that is, the interaction diameter of the filament is about four times larger than the bond length *ℓ*_0_ (Fig. 1). This prevents the geometric interlocking of neighbouring filaments when they slide past each other, and thus attenuates the sliding resistance due to the surface structure of the filament arising from the bead-spring model. The second set of simulations considers *rough filaments* with ***σ*** ≃ 1 (§4); as shown below, the corrugated filament surface resists relative tangential sliding and thus qualitatively alters the collective filament dynamics.

For all simulations, we keep the filament contour length constant: we set the number of beads *N*_m_ = 40 and set a large extensional spring constant 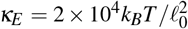 so that the filament is practically inextensible. Since the filament dynamics is sensitive to its bending rigidity, *B* we consider three values of *B* (Table 2).

**Table 2.** Amplitude 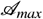 and frequency ***ω*** of oscillations of an isolated filament for three values of the activity number, *β*.

| $\beta$ | $\mathcal{A}_{\max}/\ell_0$ | $\omega \ell_0^2/D$ |
| --- | --- | --- |
| 192 | 20 | $1.4 \times 10^{-2}$ |
| 384 | 16.5 | $1.75 \times 10^{-2}$ |
| 768 | 14 | $2.1 \times 10^{-2}$ |

To mimic biological carpets such as cilia arrays, in which filaments are connected by rigid linkers or basal bodies to a wall-like substrate, we usually specify that one end of the active filament is clamped rigidly to a wall (except for Figs. 9 and 10, in which we allow the end of the filament attached to the wall to freely pivot). We initialize simulations with each filament in a straight configuration, for which the active forces are oriented toward the clamped base, causing a compressive stress along the filament.

For sufficiently large active force magnitude *f*, since the direction of *F* is aligned to the local unit vector along the arc-length of the filament and directed toward the clamped end, each filament undergoes a buckling transition and eventually nonlinear oscillations^30–32^. The follower force mechanism couples the filament configuration to the active force. Steric interactions between neighboring filaments significantly alter filament orientations and thereby the active-follower forces. Thus, filaments within a carpet undergo different dynamics than the intrinsic beating motions of isolated filaments.

### 2.4 Behavior of an isolated filament

We showed previously^29,30^ that in the case of a single clamped filament, the spatiotemporal response obtained from equations (1)–(6) depends on *β*. For *β* < *β_c_*, the filament remains nearly straight with small amplitude fluctuations in the contour due to noise. For *β* > *β_c_*, with *β_c_* = 76.2 for a noiseless system (*D* = 0), the straight filament yields to an oscillating flutter state with a well-defined frequency and amplitude.

When *β* ≫ *β_c_*, interplay between active energy injected into the oscillating filament, the elasticity of the filament, and dissipation in the ambient fluid sets the frequency of oscillation and the maximum amplitude of the oscillations. Scaling arguments then provide estimates for the frequency of oscillations^30^ 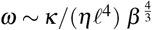 where *η* is the viscosity of the ambient fluid.

Furthermore, the oscillating filament has a well-defined amplitude whose maximum value 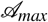 varies monotonically with *β* for the range of parameters we consider. Since the filament is clamped at one end, the lateral motion of the filament is maximal at the free end with the tip executing a figure-of-eight pattern, with amplitude 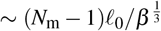. The filament tip has width *σ*, and thus moves a distance 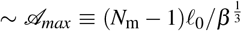. Since we ignore hydrodynamic coupling between the filaments, two filaments separated by a distance 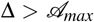 will behave predominantly as isolated filaments. The extent of steric coupling is quantified by geometric dimensionless parameters *δ* and *δ*_max_:

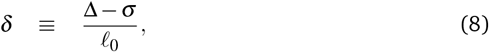

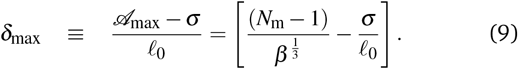

For *β* ≫ 1, we see that two filaments are closely spaced if *δ* ~ 1 and loosely spaced when *δ* ~ *δ*_max_. In Figure 1(c)-(e) we present the oscillatory dynamics of a filament in the limit *δ* ≫ *δ*_max_. For sufficiently large activity (*β* = 192) the filament undergoes regular oscillatory motion (Fig 2(a)), with a peak in the power spectrum at a frequency that depends on *β* (Figure 2(d)). Moreover, the end-segment of the filament oscillates between two maximum values, whose amplitude is denoted by 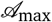.

**Fig. 2.**
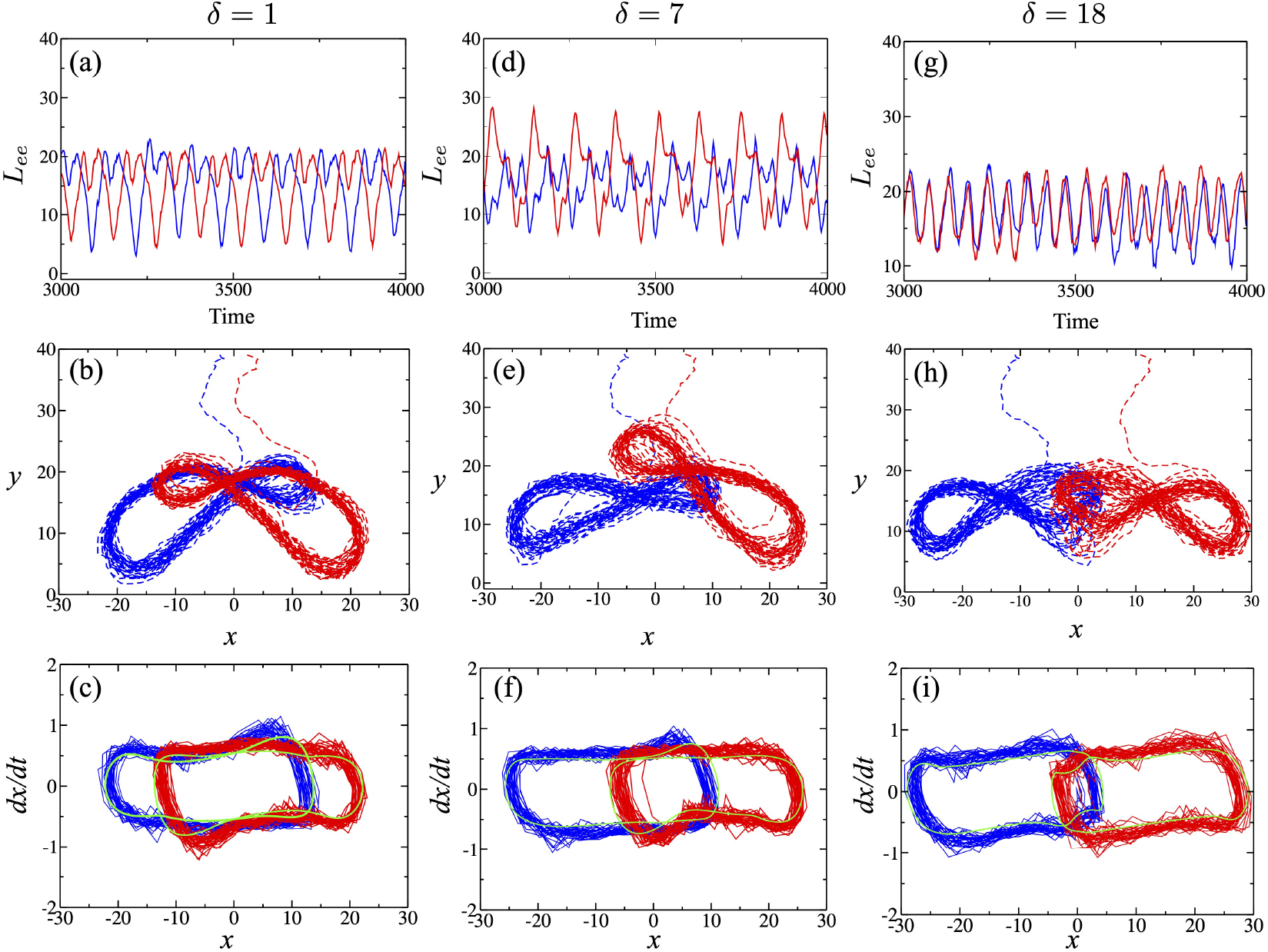
Dynamics of two clamped filaments. The length of the end-end vector *L*_ee_, the trajectory of the free end, and the *x* component of the free-end velocity as a function of the *x* position of the free end, are plotted as a function of time for a system with *N* = 2 and *β* = 384. We show (a,b,c) a relatively close pair (*δ* = 1), (d,e,f) intermediate packing (*δ* = 7) and (g,h,i) loose packing (*δ* = 18); here *δ*_max_ = 12.5. The green curves in (c,f,i) refer to the results for a simulation in the absence of thermal noise. Results are shown for smooth filaments with *σ*/*ℓ*_0_ = 4. Comparing the green and the red/blue data points, we see that in this instance, the perturbation due to noise is weak.

## 3 Small clusters of smooth filaments

An array with *N* ≫ 1 filaments may be understood as a hierarchical network, comprising of filament pairs, filaments triplets, and so on. Therefore, we first study the behaviors of a two-filament pair and a three-filament bundle to identify coordination and synchronization at small scales, followed by a large carpet (*N* = 300) to learn how these behaviors extend to larger scales. Except where mentioned otherwise we consider smooth filaments with *σ* = 4*ℓ*_0_.

### 3.1 Collective dynamics of two-filament clusters

We place two filaments with bases that are clamped and separated by a distance Δ along the *x* axis. The available space between two active filaments is then given by *δ* = (Δ − *σ*)/*ℓ*_0_. Since the isolated filament dynamics is governed by the activity number *β*, we compare the oscillatory dynamics of the filaments for three values, *β* = 768, 384, and 192. The dimensionless frequencies and maximum amplitudes in the absence of interactions 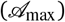 are listed for these values of *β* in Table 2. When the separation is much larger than the oscillation amplitude in the *x* direction, 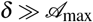, the filament interaction is zero, and we recover the oscillatory pattern for isolated filaments. Therefore, we focus on the range 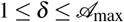. Based on the simulation results, we broadly classify the collective oscillations into three categories as follows.

#### 3.1.1 Synchronized oscillations for 1 < *δ* ≪ *δ*_max_

In Fig. 2(a-c), we summarize the dynamics of two filaments in a tightly packed configuration (*δ* = 1). We quantify the oscillatory dynamics by measuring the time-dependent end-end length *L*_ee_ of both filaments. We find that with *δ* ≃ 1, for all *β*, contact interactions qualitatively modify filament oscillations, reshaping the waveform with an additional minimum emerging in *L*_ee_(Fig. 2(a)). Moreover, there is a constant phase shift between oscillations of the two filaments. Next, we measure the trajectory of the free ends of both the filaments (Fig 2(b)). Unlike for isolated filaments, the end-segment trajectory shows an asymmetric pattern, with oscillations skewed in the direction away from the neighboring filament. Steric interactions hinder bending toward the neighbor and promote bending away from it. This mechanism also explains the appearance of the second minimum in *L*_ee_, which arises due to the occluded bending of the filament in one direction, reducing its maximal end-to-end length. In Fig. 2(c), we plot the time-derivative *dx*/*dt* of the *x* component of the end-segment position, which exhibits noisy, symmetric, closed loops for both filaments. For comparison, we show the result with thermal noise turned off in the simulations, which corresponds to smooth closed trajectories.

#### 3.1.2 Change in synchronization when 1 ≪ *δ* < *δ*_max_

Increasing the inter-filament spacing qualitatively changes the form of *L*_ee_ (Fig. 2(d)), with a significant reduction in one of the minima of the peak-to-peak oscillation compared to Fig. 2(a). Analyzing the free-end trajectory (Fig. 2(e)) shows that one of the filaments (which one is stochastic due to the left-right symmetry of the initial non-oscillating state) gets jammed under its neighbor. Thus, the oscillatory pattern of the jammed filament gets qualitatively modified while the ‘dominant’ filament oscillates with a higher amplitude that its neighbor. This behaviour is most prominent around 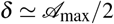. This asymmetry in interaction also affects the periodic nature of *L*_ee_, especially for the jammed filament. On the other hand, the *dx*/*dt* vs *x* dependence resembles that of the closely spaced filaments (Fig 2(f)).

For filaments with smaller activity or higher rigidity, (*β* = 194), simulations indicate that this jamming behaviour is absent (SI-Fig2). Although the interaction significantly alters the waveform in *L*_ee_ and distorts synchronization between the filaments, the effect is equal for both the filaments as neither of the filaments becomes continually jammed (see SI-§1 for details).

#### 3.1.3 Re-entrance of perfect synchronization for *δ* ≃ *δ*_max_

Increasing the inter-filament spacing to *δ* ≃ *δ*_max_ reduces the typical contact duration, and symmetry suggests that the contact occurs in the same region for either filament. Consequently, we observe statistically similar waveforms and oscillations for both filaments (Fig. 2(g)). However, the trajectories of the free ends for both filaments do overlap (Fig. 2(h)), indicating a non-vanishing contact. The periodic forcing acting on either filament combined with the activity results in a reemergence of synchronization, with this effect being more prominent for filaments with lower activity number (*β* = 192), where the *L*_ee_ waveforms of the two filaments exhibit near perfect overlap.

We can summarize the behavior of two-filament clusters as follows. With high activity number and intermediate inter-filament spacing, the two filaments exhibit different oscillatory patterns because one filament dominates and restricts the oscillations of its neighbor. In contrast, for smaller activity number the interfilament interaction leads to a similar oscillatory pattern in both the filaments with a constant phase difference (see SI§1-A and Fig-S1 for details).

### 3.2 Dynamics of three-filament clusters

We next study three filaments (*N* = 3). This geometry breaks the symmetry of the constituent filaments, since the central filament experiences steric hindrance on both sides while the end filaments each have a neighbor only on one side.

#### 3.2.1 Tightly packed filaments (*δ* ≃ 1): synchronization

All filaments interact strongly at *δ* ≃ 1. Computing the *L*_ee_ waveform and the end-point trajectory of each filament shows that the waveform is similar for both the end-filaments while it differs for the middle filament (in both amplitude and frequency, Fig 3(a)). The maximum amplitude of *L*_ee_(*t*) attained by the middle filament is roughly half that of the end-filaments and the associated frequency is almost double. The reason for this difference is evident from the end segment trajectories (Fig. 3(b)), which show that the oscillation of the middle filament is occluded by the steric hindrance due to both end-filaments. This leads to a low-amplitude, symmetric pattern for the middle filament. For the end-filaments, the oscillations are obstructed only in one direction, which leads to asymmetric patterns. This asymmetry manifests as an additional low-frequency mode in the *L*_ee_ waveform.

**Fig. 3.**
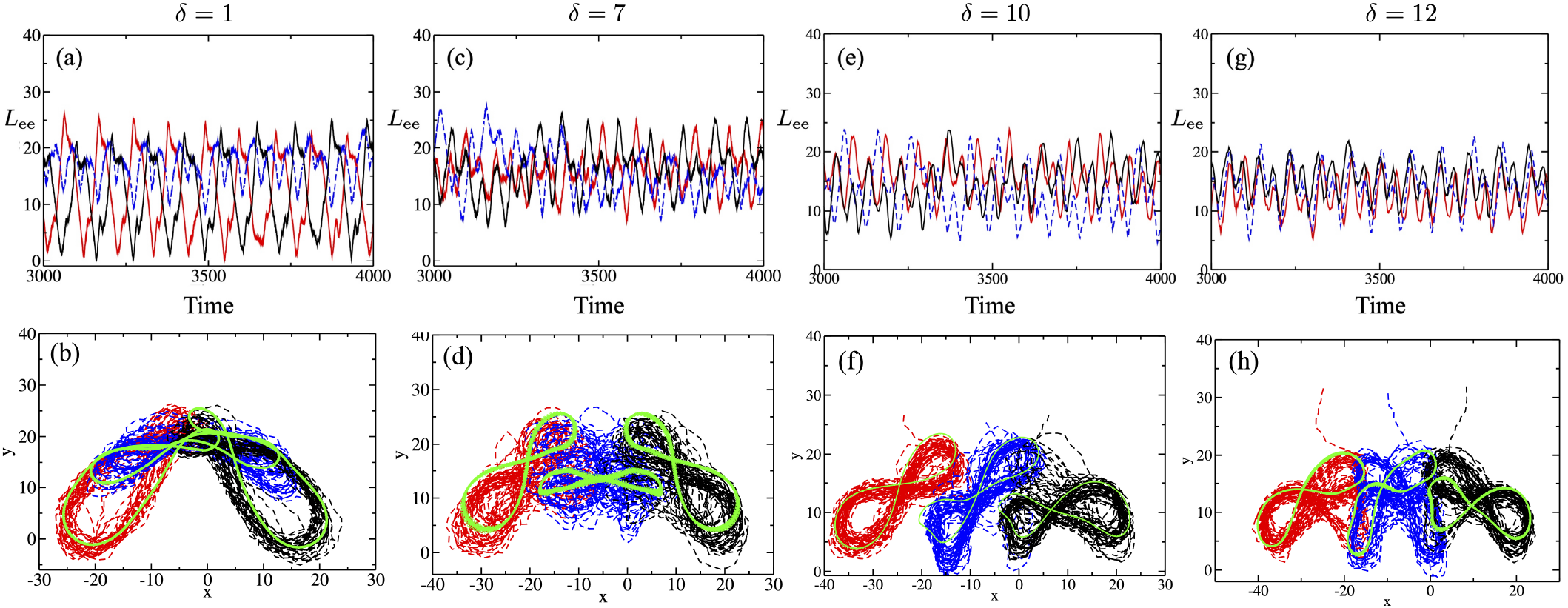
Dynamics of three clamped filaments. The time evolution of the end-end length *L*_ee_ and the end-segment trajectory are shown for inter-filament separations *δ* = 1 ((a) & (b)), *δ* = 7 ((c) & (d)), *δ* = 10 ((e) & (f)), and *δ* = 12 ((g) & (h)). All the filaments have activity number *β* = 768 so that *δ*_max_ = 10. The green curves in the second row correspond to the trajectories when noise is negligible. We note that the discreteness of the simulation scheme results in the green curves not being completely smooth. We also note the similarirties in (f) and (h) with a more pronounced asymmetry towards one of the end filaments for *δ* = 12.

#### 3.2.2 Intermediate packing (1 < *δ* < *δ*_max_): disruption and trapping

At intermediate spacing, the filaments have space to deform without contact, and we observe a disruption of regular oscillations for all the three filaments. Since the deformation depends on the filament softness (~ 1/*β*), it is especially pronounced for soft filaments with *β* = 768, where the oscillatory pattern is highly sensitive to *δ* at this range as highlighted in Fig 3(c)-(h).

At *δ* = 7, the *L*_ee_(*t*) time series shown in Fig. 3(c) shows neither regular oscillations nor synchronization, and the end-segment trajectory does not display a clear pattern (Fig. 3(b)), especially for the middle filament.

We observe a similar trend in the *dx*/*dt* vs *x* pattern at this spacing (SI§1-B). The regular oscillation is recovered when *δ* = 1Q, while the end-point trajectories of all three filaments are asymmetric but similar (Fig 3(f)). However, for *δ* = 12 (Fig 3(h)), the end-segment trajectory of the middle filament is qualitatively different compared to the end filaments. While the end-filament oscillation switches from symmetric to asymmetric patterns and back, the centre filament always oscillates asymmetrically. The direction of this asymmetry switches over time, thus resulting in an overall symmetric, butterfly-like pattern over a large time (Fig 3(g-h)).

However for stiffer filaments with *β* = 384 and 192 (SI§1-B), we do not observe such a disruption in oscillations as for soft filaments. In this case, the middle filament is trapped either below or the end filaments, restricting its oscillatory amplitude without disrupting the regular oscillations. When the separation is further increased to *δ* ≃ *δ*_max_, the filaments do not interact except for the maximally bent (minimum *L*_ee_) configurations. At this separation, we observe a reemergence of synchronized oscillations in all the filaments for all values of *β* (SI§1-C).

### 3.3 Going from *N* ~ *O*(1) to *N* ≫ 1: Anticipating the effect of contact forces

To anticipate how this time-dependent nature of the steric interactions will effect the collective behavior of *N* ≫ 1 filaments, we measure the components *f_x_* and *f_y_* and magnitude |*f*| of the *contact* forces,

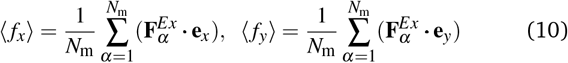

derived from the pairwise WCA potential, acting on the middle filament as a function of time for the soft filament with *β* = 384 (Fig. 4). For small basal separation (*δ* = 1), the middle filament is always in contact with the neighboring filaments and 〈*f_x_*〉 exhibits regular, albeit noisy, oscillations (Fig. 4(i)). When the basal distance is increased *δ* ≃ 10, the periodicity in 〈*f_x_*〉 weakens and the pattern is more noisy (Fig 4(ii)), which is consistent with the observed destruction of regular oscillations. At large basal distances (*δ* ≃ 17) the filament interacts with its neighbors only for a short time during the oscillation cycle, which manifests as regular pulses in 〈*f_x_*〉 pattern (Fig 4 (iii)). Such periodic pulses lead to a highly synchronized response over this range of distances.

**Fig. 4.**
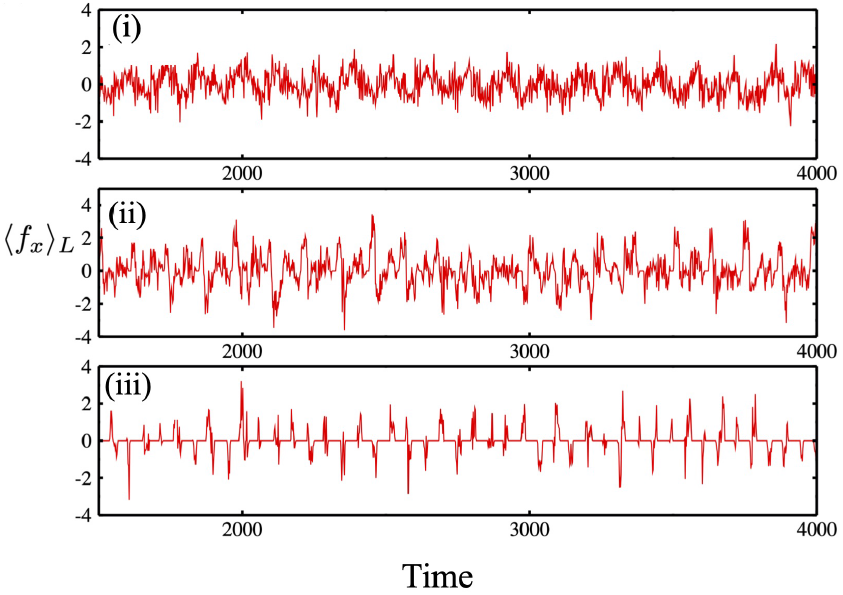
The *x* component of the mean contact force that acts on the middle filament in a three-filament cluster. (a) Here *β* = 384 (i) *δ* = 1, (ii) *δ* = 8 and (iii) *δ* = 17 (*δ*max ≃ 16.5). Plots for *β* = 192 are qualitatively similar.

## 4 Periodic array of smooth filaments

We now consider a larger system with *N* = 300 filaments arranged on a one-dimensional lattice. As above, we consider smooth filaments with uniform spacing *δ*. We apply periodic boundary conditions in the *x* direction such that the periodic images of the end filaments (1st and 300th) are also separated by *δ*, so that in the absence of spontaneous symmetry breaking, all filaments are identical. We choose an intermediate filament rigidity value, with *β* = 384.

### 4.1 Tightly packed filaments (*δ* ≃ 1): Slow metachronal waves

Under tight packing, steric interactions act on each filament throughout its oscillation cycle, which leads to a high degree of inter-filament coordination (Fig 5 (a)) (see MOVIE-1 in ESM). As in the small clusters studied above, we quantify the spatiotemporal behavior of the system via the end-end length *L*_ee_ of each filament as a function of time. We plot this information in a kymograph in Fig.5 (b), where the spatial points are the basal position of each filaments. The color code indicates *L*_ee_ of each filament with basal anchoring at *x*. The kymograph (Fig 5 (b)) indicates a phase-lag synchronization in beating between filaments separated by large distances. This manifests as metachronal waves, propagating in a specific (+*x*) direction, similar to the traveling waves observed in many biological systems. Due to the high inter-filament coordination, waveforms of each filament are similar (Fig 5 (c)).

**Fig. 5.**
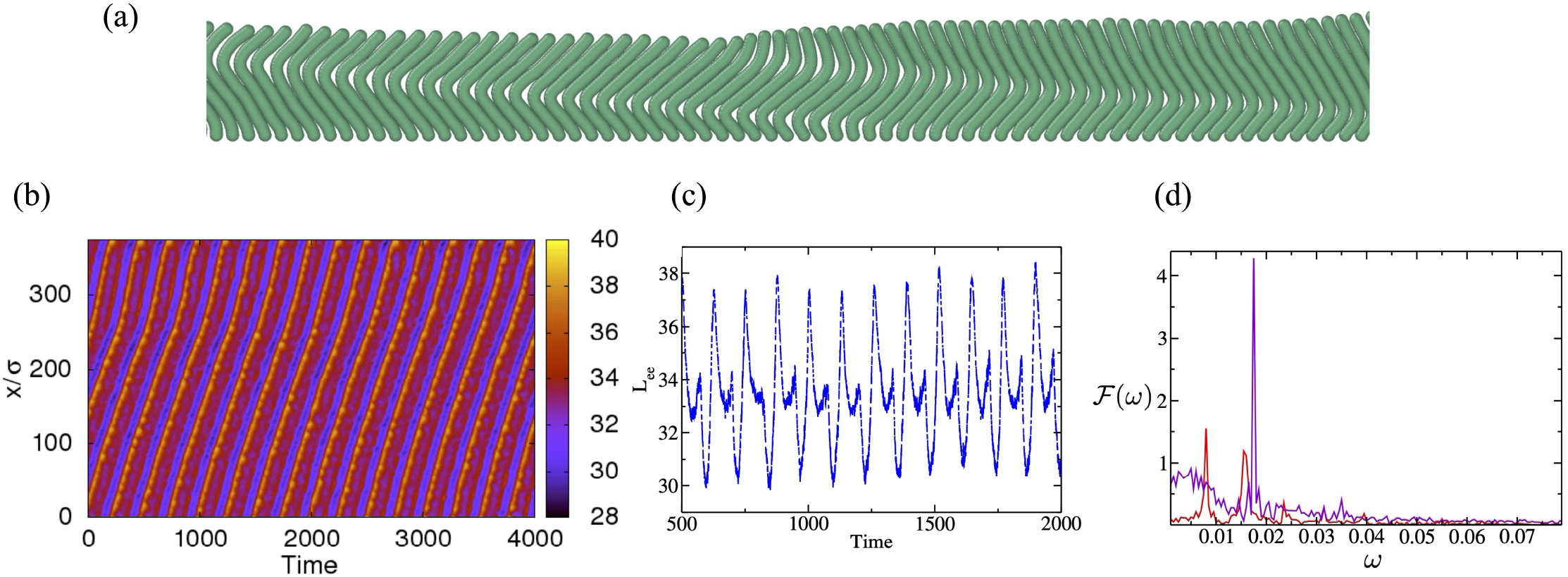
Collective dynamics of clamped filaments. (a) Snapshot of a section of *N* = 300 closely packed (*δ* = 1) clamped filaments undergoing synchronized beating at *β* = 384. Videos of corresponding simulation trajectories are shown in MOVIE-1 in the ESM. (b) Kymograph of the end-end distance *L*_ee_ of clamped filaments for *δ* = 1. The color code indicates the end-end length *L*_ee_. The 0 on the *y*-axis corresponds to the left end of the filament array. The slanted line indicates propagation of a stable waves in the +*x* direction. (c) Typical oscillatory pattern of individual filaments for *δ* = 1. The filament-filament interaction significantly reduces the filament oscillatory amplitude and frequency compared to isolated filaments. (d) Comparison of the oscillatory frequency of an individual filament inside the carpet, quantified via the Fourier transform of the end-end distance (*L*_ee_) time-series, for the tightly packed condition *δ* = 1 (red) and for isolated filaments with no inter-filament interactions *δ* ≫ *δ_max_* (purple), at *β* = 384.

However, the waveform and amplitude of *L*_ee_ are significantly different from those of an isolated filament. Fig 5(d) compares the Fourier transforms of the *L*_ee_ time-series for isolated filaments and those within the carpet, demonstrating that the steric interactions significantly reduce the oscillation frequency.

Since our results indicate that steric interaction between the filaments plays a crucial role in the emergence of cooperative oscillations, we analyze the dynamics of inter-filament forces acting on a filament due to inter-filament interactions. Fig. 6 shows kymographs of the components and magnitude of the steric forces. Since the oscillatory motion alters the local ‘contact’ of a filament in the array, the contact forces also exhibit spatiotemporal dynamics similar to Fig. 5(b). The striped pattern in Fig 6 indicates a contact propagation from the basal to the distal end of the filament. However, the periodicity in the pattern is almost double for the *F_y_* component compared to the *F_x_* component, which is specific to the filament oscillatory dynamics.

**Fig. 6.**
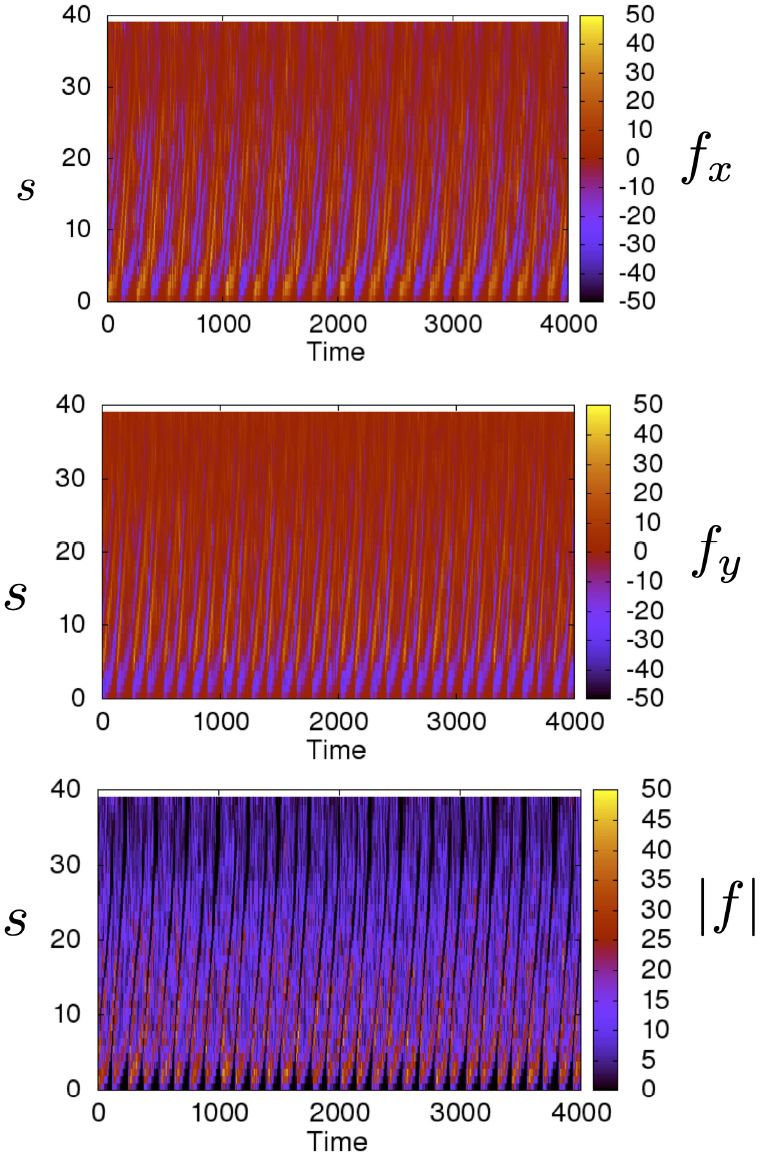
Kymographs of force components due to inter-filament repulsive interactions on sections of a filament, in a dense array of *N* = 300 smooth filaments with *δ* = 1. (a) The *x* component, (b) *y* component, and (c) magnitude |*f*|.

### 4.2 Intermediate separation: Irregular beating

Increasing the inter-filament spacing leads to disordered filament dynamics (Fig. 7 (a) and ESM MOVIE-2); the kymograph shows a lack of phase-lag synchronization or coordinated oscillations of spatially separated filaments. The lack of coordination results from irregularities in the beating patterns of individual filaments induced by interactions with their neighbors (Fig 7 (b)). Thus, the disappearance of coordinated beating at intermediate filament separations described above for *N* = 3 extends to large systems with *N* ≫ 1.

**Fig. 7.**
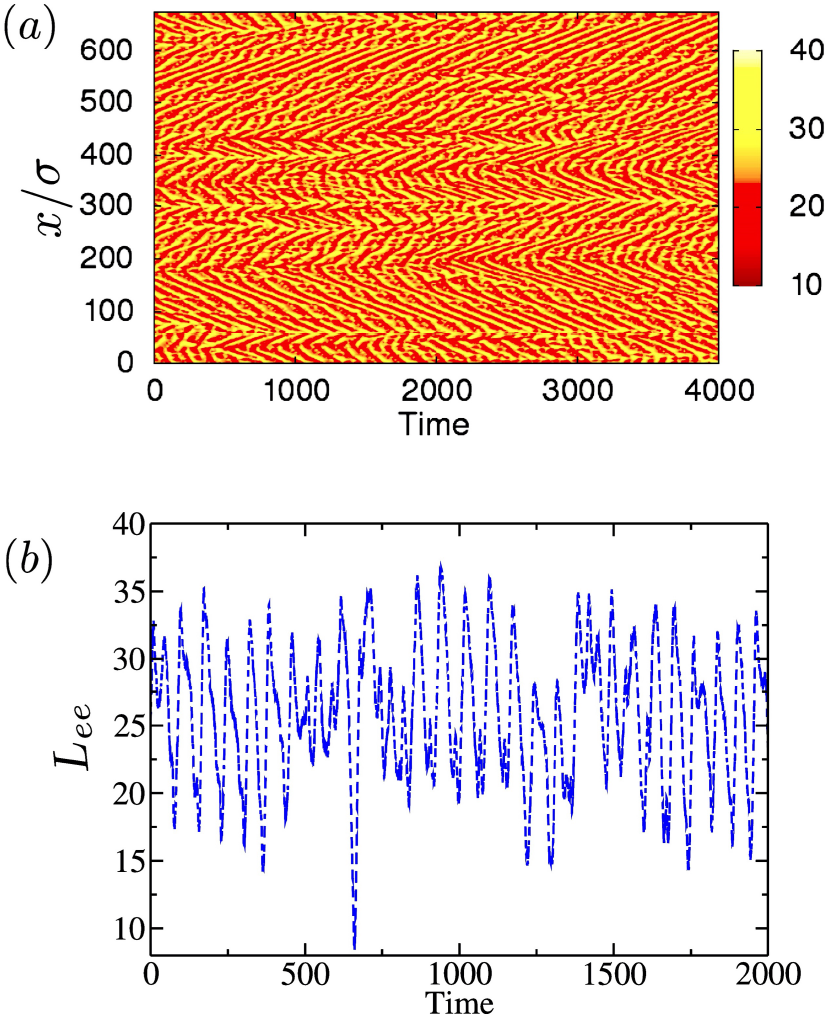
(a) Kymograph of the end-end length *L*_ee_ in system of *N* = 300 clamped filaments for the spacing parameter *δ* = 5 with *β* = 384. Videos of corresponding simulation trajectories are shown in MOVIE-2 in the ESM. The 0 on the *y*-axis corresponds to the left end of the filament array. The disordered pattern in the kymograph indicates a lack of synchronization in filament oscillations. (b) Typical waveform of *L*_ee_ of an individual filament from the same arrangement, indicating the disorder in oscillations.

### 4.3 Large separation: Emergence of fast metachronal waves

When the inter-filament spacing is further increased (*δ* > *δ*_max_/2), the contact interaction becomes ‘pulse’-like and the individual filaments beat with a higher frequency, close to that of an isolated filament. Interestingly, we observe the reemergence of waves at these large separations (Fig 8 and ESM MOVIE-2). However, the wave propagation is qualitatively different than observed for tightly packed filaments, where filaments are in continuous contact with their neighbors. At large separations, the filaments which are initially oscillating independently, coordinate their oscillatory phase through the ‘pulse’-like interactions. This results in nucleation of independent waves moving in either directions, at different regions in the array of filaments. Two oppositely moving waves meet at a ‘node’ where they annihilate (c.f Fig 8 (a)), leading to a saw-tooth pattern in the kymograph. Also, the speed of wave propagation, which is closely linked to the individual filament beating frequency, is higher compared to the tightly packed filaments.

**Fig. 8.**
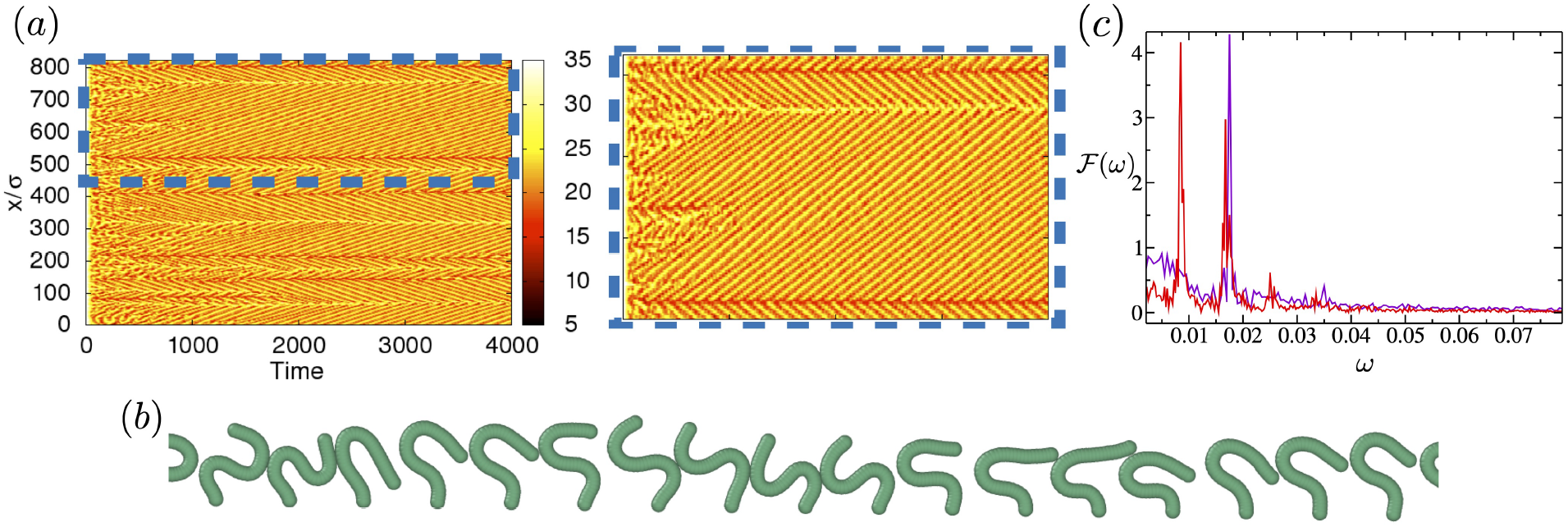
Collective dynamics of sparsely packed filaments. (a) Kymograph of the end-end length *L*_ee_ in a system of *N* = 300 clamped filaments for the spacing parameter *δ* = 11 with *β* = 384. Videos of corresponding simulation trajectories are shown in MOVIE-3 in the ESM. The 0 on the *y*-axis corresponds to the left end of the filament array. The thin, slanted patterns correspond to fast-moving waves translating in both the directions. A blown-up version of the kymograph is shown on the right. (b) Snapshot of a section of filament array, indicating a phase-lag synchronization. (c) Individual filament oscillation frequencies in a sparsely packed carpet *δ* = 11 (red) and for isolated filaments *δ* ≫ *δ_max_*.

A closer examination of the configuration (Fig 8 (b) and MOVIE-3) indicates that the filaments exhibit a phase-lagged synchronization, with a much larger phase difference compared to *δ* ≃ 1. Analysis of the frequency spectrum of *L*_ee_ oscillations identifies multiple harmonics in the oscillation waveform (Fig. 8(c)). However, the oscillation frequency of individual filaments at this separation closely matches with that of an isolated filament (Fig. 8(c)).

## 5 Periodic array of rough filaments

The previous section discusses the collective dynamics of active filaments for which the individual beads have an effective interaction diameter ***σ*** = 4*ℓ*_0_ that is larger than the equilibrium separation between neighboring beads b_0_. This arrangement ensures relatively low resistance to tangential sliding between adjacent filaments in tightly packed configurations and mimics steric interactions between brush-grafted filaments as in the mucociliary tract^47^. In this section, we discuss filaments in which the effective interaction diameter is comparable to the equilibrium inter-bead distance (***σ*** ≈ *ℓ*_0_), resulting in large gradients of the excluded-volume potential between adjacent filaments and mimicking filaments with corrugated micro-scale roughness. Effectively, beads in neighboring filaments interlock as they move, resulting in higher effective friction coefficients and significantly reducing their tangential velocities.

Additionally, we explore the role of the geometric constraint at the base in sustaining and stabilizing oscillations. Surprisingly, relaxing the hard clamped boundary condition by the softer pivot-type condition that allows for rotation leads to a new pattern - stable actively jammed structures.

### 5.1 Rough filaments with clamped bases

Fig 9 and (ESM-MOVIE-4) present the collective dynamics of *N* = 300 clamped active rough filaments. To highlight the effect of inter-filament interactions, we focus on tight packing with *δ* = 1. The activity parameter is *β* = 384. As in the case of smooth filaments, excluded volume interactions alter the phase of oscillation of individual filaments (in the array), leading to collective oscillatory patterns (Fig 9(a)). However, the patterns qualitatively differ from those exhibited by smooth filaments at *δ* = 1 (Fig. 5(b)). Instead of forming long-ranged metachronal waves that travel across the entire array, the interlocking of neighboring rough filaments results in clusters of synchronously oscillating filaments with negligible phase differences among filaments within a cluster. These clusters are separated by smaller regions of filaments that oscillate with a constant phase shift, forming short-ranged metachronal waves.

**Fig. 9.**
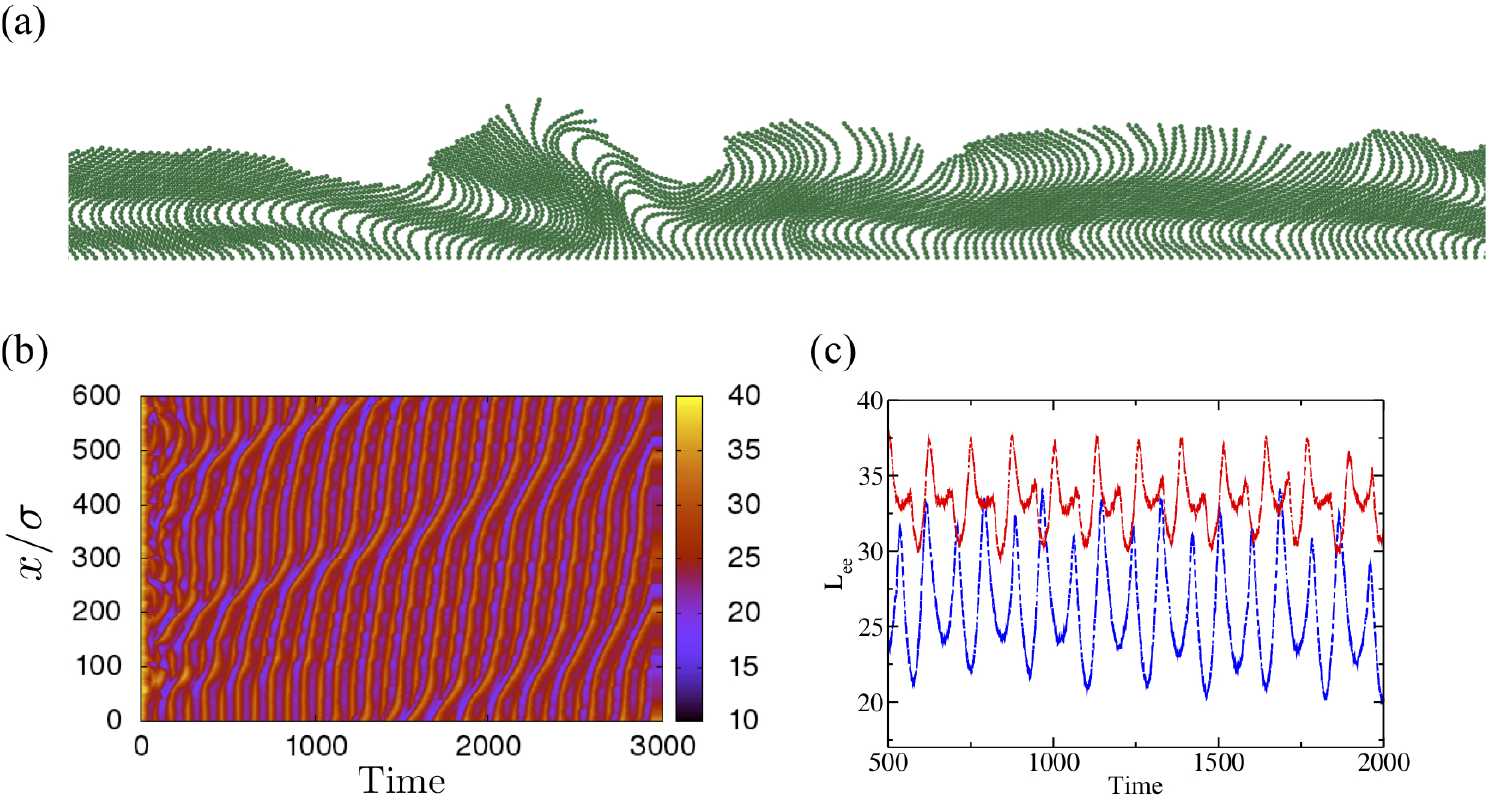
Collective dynamics of *rough* filaments (*σ*/*ℓ*_0_ = 1), with *β* = 384 and clamped at the base. (a) Typical configuration for tight packing (*δ* = 1), exhibiting regions with synchronized oscillations. (b) Kymograph of the end-end distances of the filaments. Vertically aligned stripes indicate synchronized oscillations. (c) Typical oscillatory pattern of an individual, rough filament at *δ* = 1 (red). The oscillatory pattern qualitatively differs from that observed for smooth filaments with *σ*/*ℓ*_0_ = 4 (blue).

The kymograph in Fig 9(b) illustrates this behavior, and indicates a complex collective dynamics of the filaments. The vertical stripes in the kymograph indicate groups of filaments with synchronized oscillations, while the curved regions in the stripes correspond to shifting in the location of synchronized clusters along the array. Fig. 9(c) shows the typical oscillatory pattern of individual filaments via their end-end length, *L*_ee_, which reveals the modification in oscillatory pattern of individual filaments due to crowding.

### 5.2 Rough filaments with a pivoted bases

We now consider filaments with a pivoted boundary condition at their bases, meaning rotation about the anchoring point is not energetically penalized. Our previous work showed that individual filaments with pivoted boundary conditions undergo rotational motion with a constant frequency (see^29,30,37^). Here, we examine how inter-filament interactions change this behavior by simulating an array of such filaments at *δ* = 1 (*N* = 300) and *δ* = 0.3 (*N* = 600) keeping the domain size the same. As before periodic boundary conditions are applied to the lateral ends. Note that since we do not account for excluded volumes between the filaments and anchoring surface in our simulations, the pivoted boundary condition would enable smooth filaments to slide past each other and point downward. However, for rough filaments, sliding is sufficiently restricted at small separations that this inversion does not occur. We therefore focus on rough filaments in the following.

As shown in Fig 10(a) and ESM-Movie5, we observe jammed, static clusters of filaments, separated by un-jammed filaments that oscillate. Since the net force inside a static structure must be zero, each jammed cluster has a symmetric shape; furthermore, the static clusters form at regular spatial intervals in the array. Fig 10(b) shows a kymograph of the filament end-end length for *δ* = 1, with the yellow horizontal stripes corresponding to static clusters and the slanted patterns corresponding to the small regions of oscillating filaments in between static clusters. Fig 10(c) (blue dashed line) shows the typical oscillatory pattern of the un-jammed filaments, which is similar to that of filaments with clamped boundary conditions at roughly similar *δ* (*δ* = 1.3 for the pivoting case and *δ* = 1 for the clamped case). Fig 10(c) (red solid line) also shows that the end-end length *L*_ee_ is invariant in time for filaments inside jammed clusters.

**Fig. 10.**
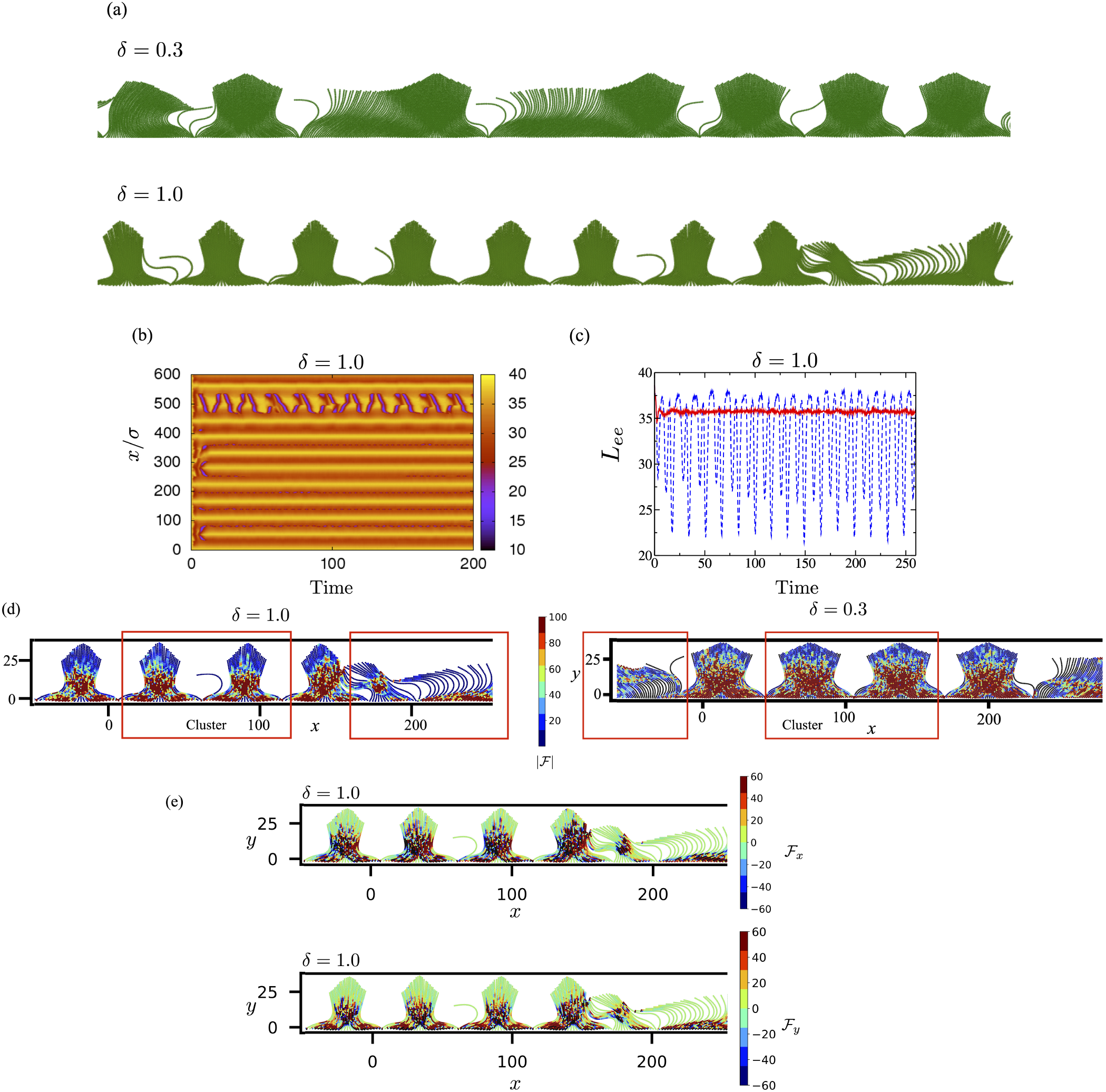
Collective dynamics of *rough* filaments with *pivoted* boundary conditions at the filament bases. (a) Typical configuration for packing densities, *δ* = 0.3 with *N* = 600 filaments and *δ* = 1.0 with *N* = 300 filaments. In both cases the filaments form jammed, static clusters, interspersed among groups of oscillating filaments. Note that periodic boundary conditions are imposed at lateral boundaries. (b) Kymograph of the end-end distance of the filaments for the *δ* = 1.0 case. Horizontal stripes indicate the static clusters. (c) End-end length of a dynamic filament with *δ* = 1, which oscillates between two static clusters (blue) and a static filament (red). (d) A map of the magnitude of total contact forces 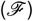 on each monomer comprising the rough filaments measured from the WCA interaction potential, for a configuration with pivoted boundary conditions for the two values of *δ* as in (a) and *β* = 384. (e) (Top) The *x* component and, (Bottom) the *y* component of the total contact forces on each bead for the configuration with *δ* = 1. Note that we show most but not all of the array; (d) may be compared to (a) to identify the part of the array shown. Periodic boundary conditions are applied such that the first and the 300th (for *δ* = 1.0) or the 600th (for *δ* = 0.3) filament are neighbors.

To understand the mechanism that drives rough filaments with pivoted boundary conditions to form jammed clusters, we analyze the inter-filament forces within jammed clusters. Fig 10(d) maps the net magnitude of the excluded-volume force (|*F*^Ex^|, eq. 5) on each bead within an arbitrarily chosen configuration. The map indicates that the interaction force is largest near the middle of the jammed cluster, where cluster undergoes maximum compression due to the active forces. In Fig 10(e)&(f), we separately analyze the *x* and *y* component of the forces respectively. We observe that the *x* component of the contact force is marginally higher compared to the *y* component, as the compression due to the outer filament acts mainly along the *x* direction. This is reminiscent of stresses borne by an arch - the distribution of compressive forces suggests that filaments can relax and unravel only by further compression given the direction of the active force thus vertically stabilizing the cluster. Lateral stabilization comes from the momentum impulses imparted to a cluster along the *x*-direction as unjammed oscillating filaments fit against the edge. Finally, there is also a geometric component due to the connected bead filament. Closer examination of the arrangement of active beads within the jammed cluster shows a nearly hexagonal packed structure that also resists sliding of beads strongly. Both these are signatures of roughness playing a dominant role.

Beyond *δ* = 2.0, we find that the clusters are very sparse since the filaments have more space in between and can rotate past each other. This is an artefact due to the lack of an actual physical barrier preventing filaments from completely sliding and moving around the pivot.

## 6 Conclusions and outlook

We have shown that purely short-ranged contact interactions are sufficient to drive coordinated beating among large arrays of active filaments, in which individual filaments beat due to compressive elastic instabilities. Moreover, such filament arrays exhibit a rich panoply of emergent behaviors, depending on the interfilament spacing, the many-body nature of the filament-filament interaction, and how filaments are attached to a surface.

Of particular interest, large arrays of smooth, tightly packed filaments exhibit highly coordinated oscillations that manifest as propagating metachronal waves. Coordination and hence metachronal waves diminish as the inter-filament spacing increases, but then reemerge at large inter-filament separations on the order of (but less than) the oscillation amplitude. Notably, the form of the metachronal waves is qualitatively different at small and large inter-filament spacing.

To understand the origin of the spatiotemporal patterns and stable states, we have systematically studied the dynamics of small clusters containing two or three filaments in addition to the large arrays. In the small tightly packed clusters of smooth filaments, coordination results in highly synchronized oscillations. Analogous to the large arrays, synchronization decreases with increasing inter-filament spacing but then reemerges at spacing comparable to the oscillation amplitude. The form of the metachronal waves in large arrays can be understood from the changes in amplitude and waveform exhibited by the small clusters at different spacing.

We also find that the nature of spatiotemporal patterns and type of stable state qualitatively differ depending on whether the filament-filament interaction is smooth or rough (corrugated) and how the filament is attached at its base. Rough filaments interlock with their neighbors at tight packing, which inhibits filament sliding motions. For rough filaments that are clamped at their base, this results in finite-size highly synchronized clusters, separated by regions of filaments undergoing asynchronous meeting. In contrast, rough filaments that freely pivot at their base form finite size *static* clusters with a size and shape that depends on the control parameters.

Two possible avenues for further work are evident. First, our results provide the foundation to study spatiotemporal patterns in active filament systems with full hydrodynamic interactions, using for instance the multi-particle collision (MPC) algorithm (e.g., Appendix in^30^). Second, our results suggest a route to understanding synchronization and collective behavior using reduced dimensional models. Current studies, focused on interactions between rotating colloids using extensions of the Kuramoto theory^39,40^, can perhaps be extended to studies of synchronization between arrays of oscillating elastic filaments. The numerical results presented here demonstrate that propagation of metachronal waves in filament arrays can arise purely via short-ranged contact interactions. While the present study is limited to a specific model for the self-regulated beating dynamics of the constituent filaments, most mechanisms that generate stable, self-regulated beating motions require coupling between the internal active force and the filament. Thus, the scope of our prediction extends beyond the particular mechanism (follower force) studied here, and can be tested in other classes of models or model systems, including those that mimic autonomous beating in cilia and flagella.

Finally, our computational model that mimics mucociliary waves can be combined with advanced numerical techniques such as Multi-Particle Collision (MPC) that allow for hydrodynamic effects or supplemented with high-resolution Galerkin methods that can treat viscoelastic interactions between moving filaments. These extensions will allow us to study the transport and capture of small particles by the bed^48^, or investigate the role of viscoelasticity^49^ in mediating inter-filament interactions in addition to steric effects explored in this paper. Viscoelastic effects introduce fluid relaxation time scales and also a means to temporarily store energy. Such simulations would be interesting, and especially relevant to understanding the effect of changes in mucus rheology on the normal functioning of cilia.

## Acknowledgements

We acknowledge supported from SERB, India via the grants SB/S2/RJN-051/2015 and SERB/F/10192/2017-2018 (RC), the UC Senate core grant (AG), the Brandeis Center for Bioinspired Soft Materials, an NSF MRSEC (DMR-1420382 and DMR-2011846) and NSF DMR-1855914 (MFH). We also acknowledge computational support from the SpaceTime2 HPC facility at IIT Bombay, NSF XSEDE computing resources (Stampede) and the Brandeis HPCC which is partially supported by DMR-1420382.

## Author Contributions

RC, AG, LM and MH together conceived the study. RC and AG designed the computational model, performed the simulations and analyzed the results. All authors contributed to the writing of the manuscript.

## Data availability

Correspondence should be addressed to RC and AG.

