## Supplementary information and figures for "Synchronized oscillations, metachronal waves, and jammed clusters in sterically interacting active filament arrays"

Raghunath Chelakkot[1], Michael F. Hagan[2], L. Mahadevan[3], and Arvind Gopinath[4]
[1]*Department of Physics, Indian Institute of Technology Bombay, Mumbai 400076, India*
[2]*Martin A. Fisher School of Physics, Brandeis University, Waltham, MA, USA.*
[3]*School of Engineering and Applied Sciences, Harvard University, Cambridge, MA, USA*
[4]*Department of Bioengineering, University of California, Merced, CA, USA*

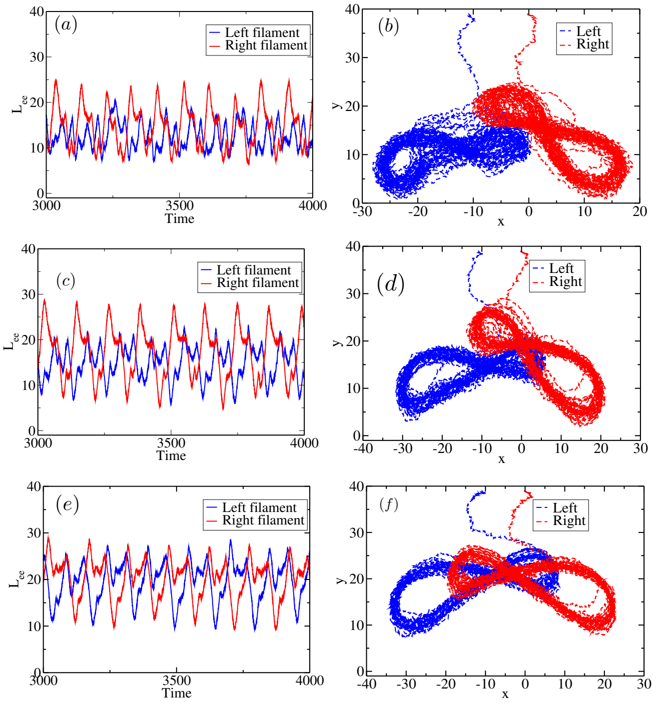


FIG. S1. The length of end-end vector $L_{ee}$ as a function of time (left panel) and the end-segment trajectory (right panel) for two filaments at intermediate separation ($\delta =$)

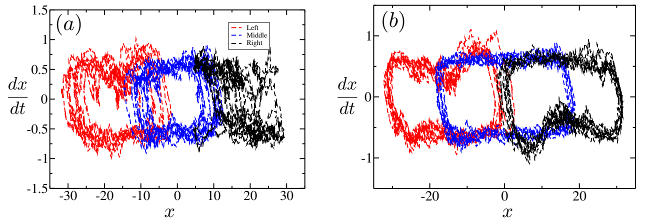


FIG. S2. Plotting the end-segment velocity component, $dx/dt$ vs. $x$ for all the three filaments at intermediate separation, $\delta = 7$. (a) for soft filament with $\beta = 768$ and (b) stiffer filament with $\beta = 384$.

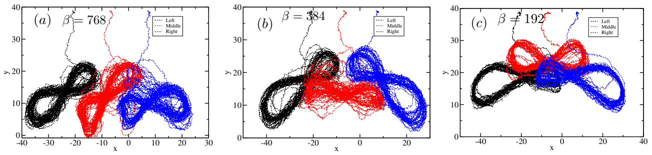


FIG. S3. The end-segment trajectory of three filament system, at intermediate separation ($\delta = 10$). The collective oscillatory pattern is different for different filament stiffness, (a) $\beta = 768$ (b) $\beta = 384$, and (c) $\beta = 192$.

### I. TWO AND THREE FILAMENT SYSTEMS AT INTERMEDIATE SEPARATION

#### A. Two filaments

At intermediate separation, the detailed dynamics of both the filaments depends on the activity number $\beta$. The filament deformation is more in the case of soft filaments ($\beta = 768$) as evident from the irregularity in their trajectories (Fig. S1(a)-(b)). Also, the difference in trajectories between both the filament indicates that one of the filaments is trapped under the other. This type of trapping also occurs for a stiffer filament with $\beta = 384$ (Fig S1(c)-(d)), although the noise in oscillations are relatively less. However the stiffest pair of filaments ($\beta = 192$) do not display trapping, since the deformation is relatively lesser in this case.

#### B. Three filaments

In an array of three filaments, the collective oscillatory pattern depends on the filament softness (via the activity number $\beta$). Similar to the two-filament system, the soft filaments ($\beta = 768$) show maximum deformation and the oscillatory patterns are different for all the three filaments (Fig S3(a)). However, for an array of stiffer filaments ($\beta 384, 192$) the oscillatory pattern of the middle

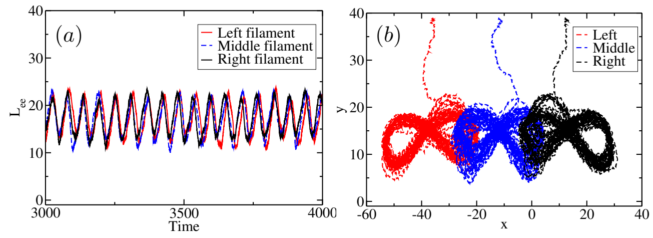


FIG. S4. (a) The end-end length $L_{ee}$ time-series and (b) the end-segment trajectory of three filaments with $\beta = 192$ and separation, $\delta = 19$ ($\delta_{\max} = 17.5$)

.

filament is different from both the end-filaments.

#### C. Three filaments - large separation

At large separation, $\delta \gtrsim \delta_{\max}$ the neighboring filaments do not interact continuously, hence the deformation caused by it is minimal. At this separation, we observe a reemergence of synhchronization between the filaments. This synchronization at large separation is particularly strong for stiffer filaments (Fig S4).
